# Red harvester ant nests impact soil compaction but not temperature in peri-urban habitats of the Lower Rio Grande Valley

**DOI:** 10.1101/2024.05.11.593703

**Authors:** Geovanni Hernandez, Hannah J. Penn, Richard Cano, Lilly V. Elliott-Vidaurri, Robin A. Choudhury

**Author notes:** Correspondence: Robin Choudhury.

## Abstract

Ants are vital ecosystem engineers that can influence soil properties, subsequent soil processes, and associated biota via underground nest construction. Harvester ants consume seeds and are often found in arid areas, frequently altering soil chemistry and bulk density of the soils in and around their nest sites. Many species of harvester ants also intentionally remove vegetation around nest openings, creating cones or discs of bare soil that may further alter soil temperatures. However, much of the work investigating the impacts of harvester ants on soil properties has occurred in shrubland and grassland settings rather than suburban environments. We aimed to determine if *Pogonomyrmex barbatus* (Smith) (Hymenoptera: Formicidae) nests in a suburban habitat in the Lower Rio Grande Valley in Texas similarly altered soil properties. First, we measured active nest disc size to determine changes and colony persistence. Then we assessed soil compaction and surface temperature along a gradient centered on the disc. We found that disc size did not increase throughout the two-year observation period and that nests with smaller discs were less likely to persist between years. While we did not observe any changes in surface temperature across the gradient, we found a significant increase in soil compaction with greater distance from the center of the disc. These data indicate that increased nest size increases the extent of soil impacted. The impacts of nests reducing soil compaction, particularly within a suburban landscape with precipitation run-off issues and a highly disturbed plant community should be addressed in future studies.

## 1. INTRODUCTION

Maintaining soil structure is critical for ecosystem services such as nutrient cycling, habitat provisioning, and water retention (Adhikari & Hartemink, 2016). Soil properties in certain environments may depend, in part, on invertebrate ecosystem engineers like earthworms, termites, and ants to create and maintain habitats (Jones et al., 1994). According to Jones et al. (1994), these ecosystem engineers both directly and indirectly alter available resources that can be used by other species by changing biotic and abiotic properties of the soil. Among other mechanisms, this can be achieved by soil invertebrates through bioturbation, the biological process of excavating, mixing, and inverting soils. Bioturbation alters soil processes such as the turnover of soil, which in turn alters organominerals, changes soil structures, and introduces new microorganisms (Jouquet et al., 2006). The presence of these ecosystem engineers, particularly those that are nest-building (e.g. ants and termites), has been shown to increase plant growth and crop yields by reducing soil compaction which then increases water infiltration and root development (De Bruyn & Conacher, 1990; Evans et al., 2011).

Ants are particularly important ecosystem engineers in soil due to their ubiquity throughout many environments and their high levels of activity, with harvester ants playing a critical role throughout global arid ecosystems (Del Toro et al., 2012; MacMahon et al., 2000a; Uhey & Hofstetter, 2022). Harvester ants are a multi-genus group of ants (including *Messor, Pogonomyrmex*, and *Veromessor*) known for preferentially feeding on seeds (Uhey & Hofstetter, 2022). This feeding alters plant communities via seed predation and dispersal as well as removal of vegetation around the nest to make a disc or cone (De Almeida, Mesléard, et al., 2020). This bare area may alter soil temperatures as these structures appear to be optimized for colony temperature requirements, particularly in cone-making *Pogonomyrmex* species (MacKay & MacKay, 1985). For instance, *Pogonomyrmex occidentalis* (Cresson) (Hymenoptera: Formicidae) bare nest cones increase internal nest temperatures via light interception (Cole, 1994) and may provide this species with extended foraging times during cooler times of the day and periods of the year in Colorado (Bucy & Breed, 2006).

Aside from surface temperature, ant-mediated bioturbation (Tiede et al., 2017) significantly changes other soil properties (De Almeida, Mesléard, et al., 2020). Specifically, harvester ant-mediated bioturbation lowers soil bulk density and increases pore space and filtration (Dostál et al., 2005; Nkem et al., 2000; Tiede et al., 2017; Whitford, 1988). For instance, soils from *Pogonomyrmex rugosus* Emery (Hymenoptera: Formicidae) nests have a lower bulk density than the surrounding soils in an Arizona mountainous desert habitat (Whitford, 1988). Lei (2000) found similar results from comparisons of P. rugosus nest located in Nevada shrublands soils with surrounding soils – nest soils had lower moisture and bulk density and compaction but greater pore space, organic matter, and temperatures, resulting in greater water infiltration.

As harvester ant nests increase in size, which is generally correlated with colony age, the amount of soil being impacted by the ant-induced bioturbation in the soil profile expands. This has been documented in both *Messor sanctus* Emery [Hymenoptera: Formicidae] and *Pogonomyrmex badius* (Latreille) [Hymenoptera: Formicidae]) colonies where the number of workers increases with colony age (Buhl et al., 2004; Tschinkel, 1999). With increased worker numbers, the need for more underground space also increases, resulting in greater nest depth and size (Buhl et al., 2004; Tschinkel, 1999). In harvester ant species that make discs or cones (MacMahon et al., 2000b), the disc/cone size also increases with colony age and size (Carlson & Whitford, 1991; Gordon & Kulig, 1996; Wiernasz & Cole, 1995). Together, these data indicate that as harvest ant colonies age and nests expand, so does the extent of soil bioturbation via both above- (via discs/cones) and below-ground processes (Carlson & Whitford, 1991).

As much of the prior work on harvester ant mediated bioturbation has been conducted in habitats with minimal anthropogenic disturbance even though harvester ants can be found in disturbed areas, we wanted to assess bioturbation impacts in a more human-mediated system. We conducted this work on red harvester ants, *Pogonomyrmex barbatus* (Smith) (Hymenoptera: Formicidae), in a peri-urban environment within the Lower Rio Grande Valley (LRGV). This species is common in Texas across many land uses including suburbs and agricultural areas and is known to make cleared discs around the nest entrance (Breene et al., 1993a; Breene et al., 1993b; Wu, 1990). Specifically, the objectives of this research were to 1) estimate changes in nest size using cleared disc diameter over time and 2) determine if the soil compaction and surface temperatures increased with greater distance from the disc center. As colony size generally increases with age, we expected that if the colony remained active (alive), nest size would increase over time (Carlson & Whitford, 1991; Gordon & Kulig, 1996; Wiernasz & Cole, 1995). Additionally, we expected soils closer to the center of the cleared disc would have lower compaction but increased surface temperatures (Bucy & Breed, 2006; Cole, 1994; Lei, 2000).

## 2. Materials and Methods

Study Site. The study was carried out across the University of Texas Rio Grande Valley (UTRGV) campus in Edinburg, TX, USA (26°18’33.1”N 98°10’26.8”W). The site (∼1.5 km2) lies in the northern section of Hidalgo County, one of four counties encompassing the LRGV, a rapidly growing sub-tropical/semi-arid region of Texas (Huang & Fipps, 2006). The LRGV is situated on the Rio Grande Plain, with deep loamy soils, moderately sloped plains, and an average elevation of 34 m (Elliott-Vidaurri et al., 2023). This region has a humid climate with a temperature range of 8 to 35°C and average annual precipitation of 609 mm.

Vegetation at the nest sites primarily consisted of grasses used for lawns with some weeds (also mostly grasses) throughout. Shrubs and trees were located further from ant nests and nearer to pedestrian walkways and buildings (Elliott-Vidaurri et al., 2023). Shrub species included standard subtropical suburban ornamental plants like *Asclepias curassavica* L. (Gentianales: Apocynaceae) and *Lantana* sp. L. (Lamiales: Verbenaceae). The site contained 53 tree species, with the majority being live oak (*Quercus virginiana* Mill. [Fagales: Fagaceae]),

Texas ebony (*Ebenopsis ebano* (Berland.) Barneby & J.W.Grimes [Fabales: Fabaceae]), and honey mesquite (*Prosopis glandulosa* Torr. [Fabales: Fabaceae]) (UTRGV Office of Sustainability, 2020). Areas surrounding the study site were primarily suburban and peri-urban with intermixed row crop fields, pasture, and citrus groves (USDA National Agricultural Statistics Service, 2022).

### Nest Measurements

Active nests were initially located within the study site in 2020 and GPS coordinates of each nest recorded (Elliott-Vidaurri et al., 2023). Active nest disc diameters were recorded annually during the summers of 2020 and 2021. Nest diameters were measured from the edge of active vegetation growth, and three diameter measurements were taken for each nest at each time point. Nests across all sites were measured over the course of seven days, limiting the effects of intra-year differences between nest-site sizes.

### Soil Properties

Soil compaction and surface temperatures relative to the nest disc were recorded for 43 colonies in the summer of 2023. Measurements were taken at 60 and 120cm starting at the nest entrance proceeding both north and south of the nest entrance. Surface temperature measurements were collected using an infrared thermometer model 62-MAX (Fluke Corporation, Everett, WA, USA) and soil compaction measurements were collected using a soil compaction tester probe model 2LBB3 (Dickey-John Inc. Auburn, IL, USA). To adjust for fluctuations in surface temperature due to an increase in diurnal temperature, all measurements were standardized to the measurement at the center of the nest by dividing all temperature measurements by the temperature at the center of each nest.

### Statistical Analyses

The size of ant nests in 2020 and 2021 were compared using a paired t-test in the R statistical program language v.4.3.3 (R Core Team, 2024), comparing the nests that were recorded in both 2020 and 2021. Additionally, all nest measurements from 2020 and 2021 were compared using an unpaired t-test to account for any biases that may have been introduced from exclusion of new nests that were assessed in 2021. A logistic regression to model the effects of colony diameter on binary nest persistence (persist/not persist) was fitted using a generalized linear model with a binomial distribution and a logit link function. A receiver operating characteristic (ROC) curve was developed to assess the effectiveness of nest size as a predictor for nest persistence, and a Tjur’s coefficient of discrimination and a Youden’s J statistic were calculated to assess the overall effectiveness of the model. Tjur’s coefficient of discrimination describes the difference between the mean predicted probability of persistence and non-persistence. Youden’s J index ranges between -1 and 1, and a value of 1 would indicate a perfect predictor model that had no false positive or false negative values. The effects of distance and direction from the center of the ant nests on surface temperature and soil compaction was assessed using a linear mixed model with a Gaussian distribution with direction and colony as random effects, with random intercepts and random slopes for direction (north and south) for each colony.

## 3. Results

### Nest Measurements

Harvester ant nests were dispersed widely throughout campus. The vast majority of nests were located on the periphery of campus, away from walking paths and trees and present mostly in open field areas. The landscape of the site was actively maintained, with regular mowing of grassy areas, removal of excess leaf litter and weeds, and irrigation of some areas. The number of nests monitored for diameter in 2020 was 45 and the number of nests in 2021 was 119. The average disc diameter was 78.2 ± 31.6 cm in 2020 and 67.9 ± 31.1 cm in 2021 (Figure 1a). In general, the results of the paired t-test suggest that nest disc sizes did not significantly change over the observation periods (t = 0.677, p = 0.504). When accounting for all nests monitored in 2020 and 2021, the unpaired t-test also suggested that there was no significant difference in the size of nests between the two years (t = 0.065, p = 0.065).

**Figure 1.**
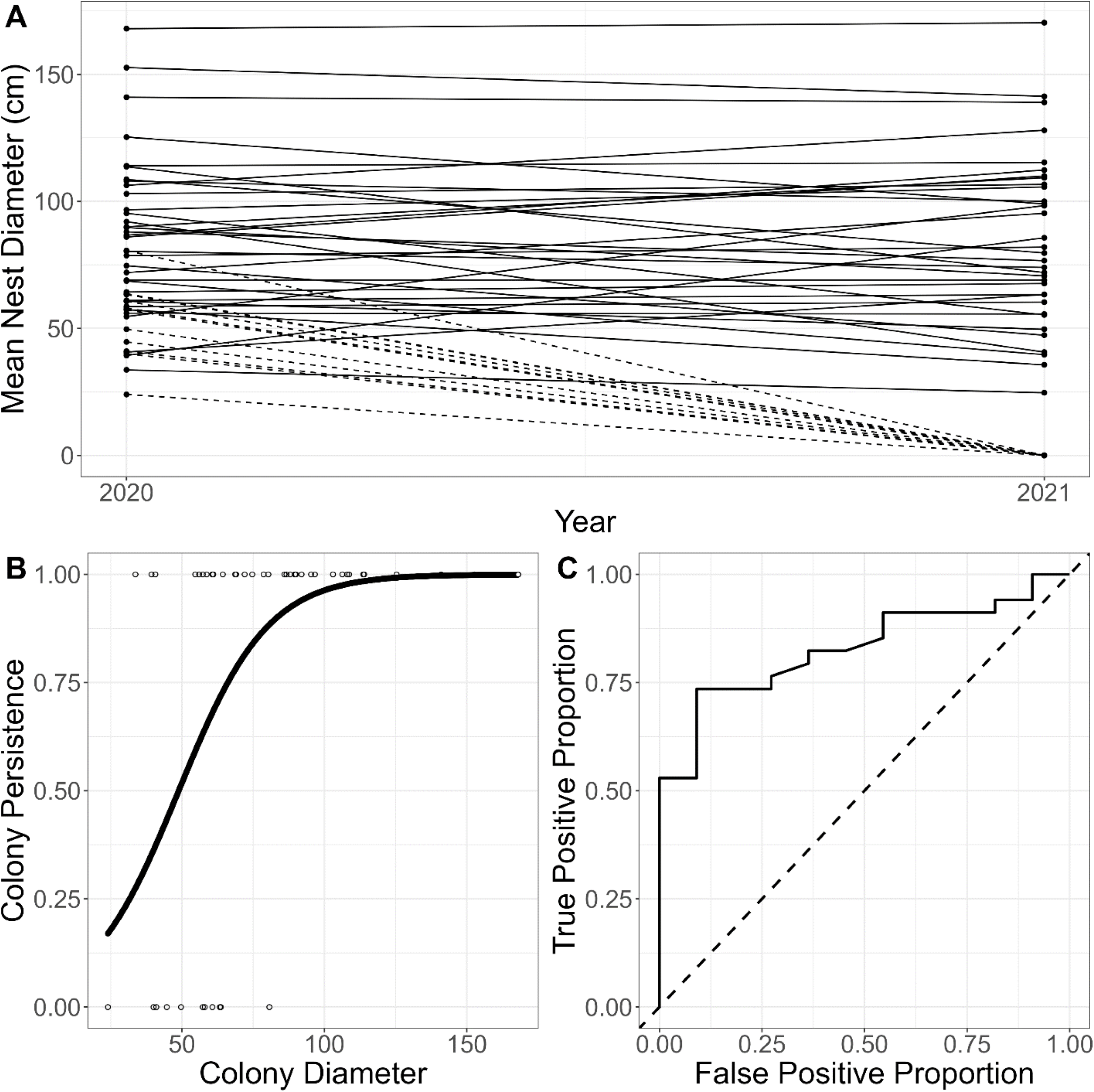
Change in size and persistence of red harvester ant nests between 2020 and 2021 (A), where the solid lines represent colonies that were seen in both years and the dotted lines represent colonies that were observed in 2020 but not 2021. The logistic regression of colony persistence by colony size (B), with open circles representing true data points and the solid black line representing the results of the logistic regression. The solid line represents the receiver operating characteristic curve of the logistic regression (C), with the dotted line representing the line of no discrimination.

### Nest Persistence

Overall, 11 nest discs that were monitored in 2020 were not present in 2021, out of a total of 45. The average 2020 nest disc diameter of nests that persisted from 2020 to 2021 was 86.3 ± 31.3 cm while the average nest disc diameter of those that did not persist was 53 ± 15.2 cm. The logistic regression of the effect of nest size on nest persistence suggests that larger nests were more likely to survive between years (p = 0.007, Figure 1b), with a log odds coefficient of 0.064. This suggests that for every centimeter increase in nest diameter, the odds of surviving to the next year increase by about 6.6%. The ROC analysis suggest that nest size is a good predictor of nest persistence, with a Tjur’s coefficient of discrimination of 0.266 and a Youden’s J index statistic of 0.644 (Figure 1c). The relatively high Tjur’s coefficient of discrimination indicated that there was a relatively large gap in the predicted probability of persistence and non-persistence in our model, and the relatively high Youden’s J index indicated that our model had relatively few false positives and false negatives.

### Soil Properties

We found that there was less soil compaction in soils closer to the center of nest discs (Figure 2). The mean compaction at 0, 60, and 120 cm from the center of the disc was 9.2, 11.4, and 12.4 kg/cm2, respectively. The results of the generalized linear model suggest that compaction significantly increased at a rate of approximately 22.9 g/cm2 for every one-centimeter distance from the center of the nest (p = 2.07e-15). There was no significant effect of sampling direction on the soil compaction in and around the disc (p = 0.154). The overall model had a substantial explanatory power (R2 = 0.79). We also found that there was no significant difference in the surface temperature relative to the distance from the center of the nest ((p = 0.458); Figure 3). The direction of measurement did not significantly affect this relationship (p = 0.933), and the overall explanatory power of the model was weak (R2 = 4.14e-3).

**Figure 2.**
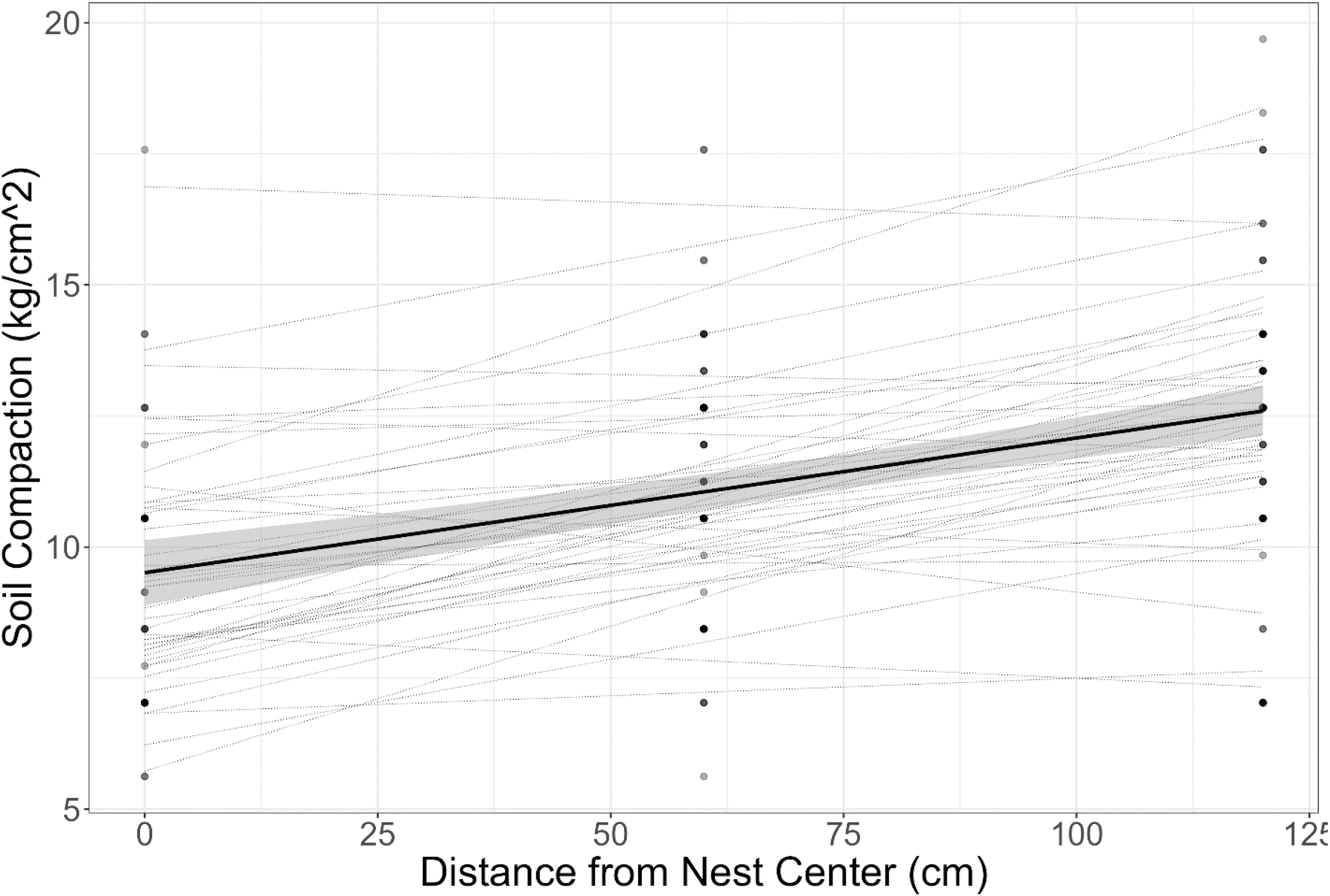
The soil compaction of a harvester ant nest when compared to distances away from the nest center. Dotted lines represent the linear regression results for each individual nest and the solid black line represents the overall linear model.

**Figure 3.**
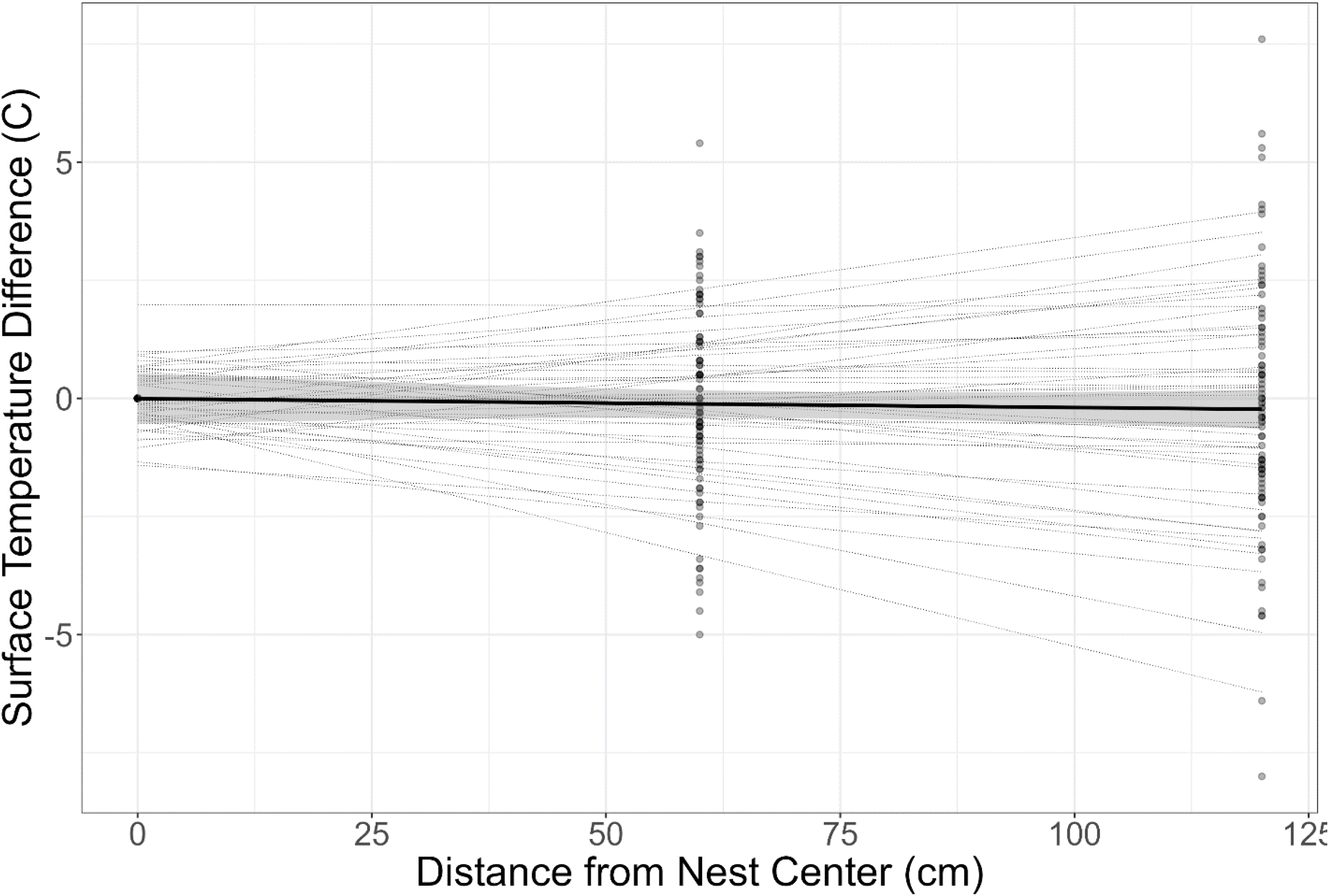
The surface temperature of a harvester ant nest when compared to distances away from the nest center. Dotted lines represent the linear regression results for each individual nest and the solid black line represents the overall linear model.

## 4. Discussion

One of the primary goals of our study was to evaluate *P. barbatus* cleared nest disc size and persistence over time. We found that *P. barbatus* nest discs did not increase in diameter over a two-year period, with nest disc size remaining steady in our peri-urban settings. These results differed from prior studies on *P. barbatus* (Gordon & Kulig, 1996) and *P. occidentalis* colonies (Wiernasz & Cole, 1995) in more natural environments. Previous studies have linked the size of harvester ant nest discs with their overall population (Tschinkel, 1999; Tschinkel, 2014). It is possible that limitations of preferred seeds in urban environments may also limit the ability of harvester ants colonies to expand worker numbers and associated nest size relative to more seed-rich natural environments (Elliott-Vidaurri et al., 2022; MacMahon et al., 2000b). It is also possible that managed landscapes in peri-urban settings may limit the nest persistence and growth.

Some variation in disc size over time may be attributable to the death and recolonization or movement of colonies from specific nest sites, as about 10% of *P. barbatus* colonies moved annually in a study conducted in New Mexico (Gordon, 1992). We found that colonies with smaller nest discs were more likely to be absent from that exact location in the second year of sampling. It is also possible that smaller nests were more likely to relocate, moving from their original site to nearby sites, but the limited number of new nests near the disappearing nests suggests that several of the nests died rather than relocated. Pinter-Wollman and Brown (2015) found that colonies of the true harvester ant (Veromessor andrei) only relocated within approximately five to ten meters from their original location. Similarly, Tschinkel (2014), who found that *P. badius* nests relocated on average 4.1m, determined that relocations were unrelated to colony size. Overall, the results suggest that smaller nests were less likely persist between years, indicating that nest size is associated with year-to-year nest survival. These results are similar to those found by Cole et al. (2022), who found there was a positive correlation between colony size and colony survival of young *P. occindentalis* colonies that were artificially transplanted into the environment. Survival may depend, in part, on food availability that impacts the health of individual ants and the overall health of the colony. Pol et al. (2022) found that nests in heavily grazed areas had fewer workers and fewer individuals per colony when compared with lightly grazed areas, and that individuals from heavily grazed areas had significantly lower fat content. The *V. andrei* nest relocations monitored by Pinter-Wollman and Brown (2015) were affected by population density and environmental factors, with more relocation at sites with lower population density and higher humidity.

The other main goal of our work was to assess impacts of red harvester ant nest bioturbation on soil compaction and surface temperatures in a peri-urban setting. Our work indicated that soil compaction increased with distance from the disc center, regardless of sampling direction. The decrease in soil compaction indicates that the soils within and nearest the nest discs may have a higher capacity for water infiltration. Lei (2000) monitored the effects of seed harvester ants on soil attributes in a desert shubland and found that ant nest soils had significantly lower soil compaction and soil moisture and significantly increased soil temperature and water infiltrability. Cammeraat et al. (2002) also found a higher infiltrability in one of the years that they monitored, although the infiltrability was reduced in warmer and drier conditions. Given this, harvester ant bioturbation may help to alleviate negative effects associated with heavily compacted peri-urban soils (Dostál et al., 2005; Tiede et al., 2017), which is particularly important in arid regions like the LRGV (Cammeraat et al., 2002; Lei, 2000). Elliott-Vidaurri et al. (2023) found that most P. barbatus nests in a peri-urban environment were within ten meters of impermeable surfaces like roads or walkways, and a similar phenomenon was observed by Uhey et al. (2021) for *P. barbatus* nests near hiking trails with high soil compaction in Arizona. One possible reason is that precipitation runoff from these less permeable and more compacted surfaces could improve the infiltration and ability of ants to excavate and establish nests in nearby soils.

Unlike compaction, surface temperatures did not differ between discs and the surrounding soils, unlike a study of *P. rugosus* nests where nests increased soil temperature (Lei, 2000). Our data are more similar to that for *V. andrei* where nest presence did not alter temperatures (Boulton et al., 2003). This may be due to the time of year or day sampling occurred, as nest cones and discs are more impactful during cooler periods (Bucy & Breed, 2006; Crist & Williams, 1999; MacKay & MacKay, 1985). One possibility for the lack of temperature difference is the effect of surrounding vegetation on surface temperatures. In a prior study, Elliott-Vidaurri et al. (2023) found that nests were unlikely to be under canopy cover, and were common in lawns. Further, the relatively sparse and low grass-cover found in many peri-urban settings does not reduce surface temperatures as much as tree-cover, especially in warm summer months (Ni et al., 2019). However, the lack of interaction between our observed discs and surface may not impact colony survival. Harris et al. (2024) found that ants from urban and rural environments had significantly different critical thermal stress temperatures, and that ants from urban environments generally thrived more in higher thermal stress when compared with rural populations. Similarly, Diamond et al. (2018) found that ant populations from some urban environments had less cold tolerance than those from rural areas, although the results differed based on the city location.

Ultimately, our study helps reveal how harvester ants shape and interact with human-altered ecosystems. We found that harvester ant colonies in peri-urban environments directly impact soil properties in similar ways observed in harvester ants located in less anthropogenically modified habitats. While the presence of harvester ant nests decreased soil compaction locally, they did not significantly alter overall surface temperatures compared with nearby grassy areas. This limited effect of nest presence on surface temperature may change over the course of a year, and future studies of *P. barbatus* in peri-urban settings can focus on long-term monitoring of nest health and their effects on the environment. Other studies have found that such ant-mediated alterations of soil properties may influence the amount of plant richness and growth (Brown et al., 2012; Whitford, 1988), which may, in turn, increase other ecosystem services (De Almeida et al., 2020b). Additionally, the presence of harvester ant nests may alter plant communities to be more similar to those in natural areas (De Almeida et al., 2020a) and increase plant reproduction and growth Farji-Brener and Werenkraut (2017). These benefits may indicate that harvester ants in peri-urban systems may also assist in restoring native plant communities as seen in grasslands De Almeida et al. (2020a).

Harvester ants are critical components within many arid ecosystems, serving as food for endangered species like the Texas horned lizard (Whiting et al., 1993). Ultimately, harvester ants are a critical component of the south Texas wildlife and their continued survival in altered settings suggests they may adapt well to human-modified conditions.

## Acknowledgements

We would like to Dr. Alice A. Wright for manuscript comments. Mention of trade names or commercial products in this publication is solely for the purpose of providing specific information and does not imply recommendation or endorsement by the U.S. Department of Agriculture. USDA is an equal opportunity provider and employer.

## Author contributions

RAC and HP contributed to the study conception and design. Material preparation and data collection were performed by RAC, GH, RC, and LVE. All analyses and figure development were performed by GH and RAC. The first draft of the manuscript was written by GH, RAC, and HP and all authors commented on previous versions of the manuscript. All authors read and approved the final manuscript.

## Data availability

The datasets and code generated during and/or analyzed during the current study are available upon request.

## References Cited

Adhikari, K. & Hartemink, A.E. (2016) Linking soils to ecosystem services—A global review. Geoderma, 262, 101–111.

Boulton, A.M., Jaffee, B.A., & Scow, K.M. (2003) Effects of a common harvester ant (Messor andrei) on richness and abundance of soil biota. Applied Soil Ecology, 23, 257–265.

Breene, R., Dean, D., & Meagher, R. (1993a) Spiders and ants of Texas citrus groves. The Florida Entomologist, 76, 168–170.

Breene, R., Meagher, R., & Dean, D. (1993b) Spiders (Araneae) and ants (Hymenoptera: Formicidae) in Texas sugarcane fields. The Florida Entomologist, 76, 645–650.

Brown, G., Scherber, C., Ramos Jr, P., & Ebrahim, E.K. (2012) The effects of harvester ant (Messor ebeninus Forel) nests on vegetation and soil properties in a desert dwarf shrub community in north-eastern Arabia. Flora-Morphology, Distribution, Functional Ecology of Plants, 207, 503–511.

Bucy, A.M. & Breed, M.D. (2006) Thermoregulatory trade-offs result from vegetation removal by a harvester ant. Ecological Entomology, 31, 423–429.

Buhl, J., Gautrais, J., Deneubourg, J.-L., & Theraulaz, G. (2004) Nest excavation in ants: group size effects on the size and structure of tunneling networks. Naturwissenschaften, 91, 602–606.

Cammeraat, L., Willott, S., Compton, S., & Incoll, L. (2002) The effects of ants’ nests on the physical, chemical and hydrological properties of a rangeland soil in semi-arid Spain. Geoderma, 105, 1–20.

Carlson, S.R. & Whitford, W.G. (1991) Ant mound influence on vegetation and soils in a semiarid mountain ecosystem. American Midland Naturalist, 125–139.

Cole, B. (1994) Nest architecture in the western harvester ant, Pogonomyrmex occidentalis (Cresson). Insectes Sociaux, 41, 401–410.

Cole, B.J., Jordan, D., LaCour-Roy, M., O’Fallon, S., Manaker, L., Ternest, J.J., Askew, M., Garey, D., & Wiernasz, D.C. (2022) The benefits of being big and diverse: early colony survival in harvester ants. Ecology, 103, e03556.

Crist, T.O. & Williams, J.A. (1999) Simulation of topographic and daily variation in colony activity of Pogonomyrmex occidentalis (Hymenoptera: Formicidae) using a soil temperature model. Environmental Entomology, 28, 659–668.

De Almeida, T., Blight, O., Mesléard, F., Bulot, A., Provost, E., & Dutoit, T. (2020a) Harvester ants as ecological engineers for Mediterranean grassland restoration: Impacts on soil and vegetation. Biological Conservation, 245, 108547.

De Almeida, T., Mesleard, F., Santonja, M., Gros, R., Dutoit, T., & Blight, O. (2020b) Above-and below-ground effects of an ecosystem engineer ant in Mediterranean dry grasslands. Proceedings of the Royal Society B, 287, 20201840.

De Bruyn, L.L. & Conacher, A.J. (1990) The role of termites and ants in soil modification-a review. Soil Research, 28, 55–93.

Del Toro, I., Ribbons, R.R., & Pelini, S.L. (2012) The little things that run the world revisited: a review of ant-mediated ecosystem services and disservices (Hymenoptera: Formicidae). Myrmecological News, 17, 133–46.

Diamond, S.E., Chick, L.D., Perez, A., Strickler, S.A., & Martin, R.A. (2018) Evolution of thermal tolerance and its fitness consequences: parallel and non-parallel responses to urban heat islands across three cities. Proceedings of the Royal Society B: Biological Sciences, 285, 20180036.

Dostál, P., Březnová, M., Kozlí;čková, V., Herben, T., & Kovář, P. (2005) Ant-induced soil modification and its effect on plant below-ground biomass. Pedobiologia, 49, 127–137.

Elliott-Vidaurri, L.V., Martinez, I., Pereira, E., Penn, H.J., & Choudhury, R.A. (2023) Tree canopy cover and elevation affect the distribution of red harvester ant nests in a peri-urban setting. Environmental Entomology, 52, 510–520.

Elliott-Vidaurri, L.V., Rivera, D., Noval, A., Choudhury, R.A., & Penn, H.J. (2022) Red Harvester ant (Pogonomyrmex barbatus F. Smith; Hymenoptera: Formicidae) preference for cover crop seeds in South Texas. Agronomy, 12, 1099.

Evans, T.A., Dawes, T.Z., Ward, P.R., & Lo, N. (2011) Ants and termites increase crop yield in a dry climate. Nature Communications, 2, 262.

Farji-Brener, A.G. & Werenkraut, V. (2017) The effects of ant nests on soil fertility and plant performance: a meta-analysis. Journal of Animal Ecology, 86, 866–877.

Gordon, D.M. (1992) Nest relocation in harvester ants. Annals of the Entomological Society of America, 85, 44–47.

Gordon, D.M. & Kulig, A.W. (1996) Founding, foraging, and fighting: colony size and the spatial distribution of harvester ant nests. Ecology, 77, 2393–2409.

Harris, B.A., Stevens, D.R., & Mathis, K.A. (2024) The effect of urbanization and temperature on thermal tolerance, foraging performance, and competition in cavity-dwelling ants. Ecology and Evolution, 14, e10923.

Huang, Y. & Fipps, G. (2006) Landsat satellite multi-spectral image classification of land cover change for GIS-based urbanization analysis in irrigation districts: evaluation in the Lower Rio Grande Valley. Texas Water Resources Institute, College Station, USA.

Jones, C.G., Lawton, J.H., & Shachak, M. (1994) Organisms as ecosystem engineers. Oikos, 373–386.

Jouquet, P., Dauber, J., Lagerlöf, J., Lavelle, P., & Lepage, M. (2006) Soil invertebrates as ecosystem engineers: intended and accidental effects on soil and feedback loops. Applied Soil Ecology, 32, 153–164.

Lei, S.A. (2000) Ecological impacts of seed harvester ants on soil attributes in a Larrea-dominated shrubland. Western North American Naturalist, 439–444.

MacKay, W. & MacKay, E. (1985) Temperature modifications of the nest of Pogonomyrmex montanus (Hymenoptera: Formicidae). The Southwestern Naturalist, 30, 307–309.

MacMahon, J.A., Mull, J.F., & Crist, T.O. (2000a) Harvester ants (Pogonomyrmex spp.): their community and ecosystem influences. Annual Review of Ecology and Systematics, 31, 265–291.

MacMahon, J.A., Mull, J.F., & Crist, T.O. (2000b) Harvester ants (Pogonomyrmex spp.): their community and ecosystem influences. Annual Review of Ecology Systematics, 31, 265–291.

Ni, J., Cheng, Y., Wang, Q., Ng, C.W.W., & Garg, A. (2019) Effects of vegetation on soil temperature and water content: Field monitoring and numerical modelling. Journal of Hydrology, 571, 494–502.

Nkem, J., De Bruyn, L.L., Grant, C., & Hulugalle, N. (2000) The impact of ant bioturbation and foraging activities on surrounding soil properties. Pedobiologia, 44, 609–621.

Pinter-Wollman, N. & Brown, M.J. (2015) Variation in nest relocation of harvester ants is affected by population density and food abundance. Behavioral Ecology, 26, 1569–1576.

Pol, R.G., Miretti, F., & Marone, L. (2022) Lower food intake due to domestic grazing reduces colony size and worsens the body condition of reproductive females of harvester ants. Journal of Insect Conservation, 26, 583–592.

R Core Team, R. (2024) R: A language and environment for statistical computing.

Tiede, Y., Schlautmann, J., Donoso, D.A., Wallis, C.I., Bendix, J., Brandl, R., & Farwig, N. (2017) Ants as indicators of environmental change and ecosystem processes. Ecological Indicators, 83, 527–537.

Tschinkel, W.R. (1999) Sociometry and sociogenesis of colonies of the harvester ant, Pogonomyrmex badius: distribution of workers, brood and seeds within the nest in relation to colony size and season. Ecological Entomology, 24, 222–237.

Tschinkel, W.R. (2014) Nest relocation and excavation in the Florida harvester ant, Pogonomyrmex badius. PLoS One, 9, e112981.

Uhey, D.A., Cummins, G.C., Rotter, M.C., Lassiter, L.S., & Whitham, T.G. (2021) Hiking trails increase abundance of harvester ant nests at Clear Creek, Arizona. Southwestern Entomologist, 46, 403–412.

Uhey, D.A. & Hofstetter, R.W. (2022) From pests to keystone species: Ecosystem influences and human perceptions of harvester ants (Pogonomyrmex, Veromessor, and Messor spp.). Annals of the Entomological Society of America, 115, 127–140.

UTRGV Office of Sustainability (2020). UTRGV 2020 Tree Campus USA Report, The Univrersity of Texas Rio Grande Valley, Office of Sustainability.

Whitford, W.G. (1988) Effects of harvester ant (Pogonomyrmex rugosus) nests on soils and a spring annual, Erodium texanum. The Southwestern Naturalist, 33, 482–485.

Whiting, M.J., Dixon, J.R., & Murray, R.C. (1993) Spatial distribution of a population of Texas horned lizards (Phrynosoma cornutum: Phrynosomatidae) relative to habitat and prey. The Southwestern Naturalist, 38, 150–154.

Wiernasz, D.C. & Cole, B.J. (1995) Spatial distribution of Pogonomyrmex occidentalis: recruitment, mortality and overdispersion. Journal of Animal Ecology, 519–527.

Wu, H. (1990) Disk clearing behavior of the red harvester ant, Pogonomyrmex barbatus Smith. Bulletin of the Institute of Zoology, Academia Sinica, 29, 153–164.

